# Maximum Likelihood Reconstruction of Ancestral Networks by Integer Linear Programming

**DOI:** 10.1101/574814

**Authors:** Vaibhav Rajan, Carl Kingsford, Xiuwei Zhang

## Abstract

**Motivation:** The study of the evolutionary history of biological networks enables deep functional understanding of various bio-molecular processes. Network growth models, such as the Duplication-Mutation with Complementarity (DMC) model, provide a principled approach to characterizing the evolution of protein-protein interactions (PPI) based on duplication and divergence. Current methods for model-based ancestral network reconstruction primarily use greedy heuristics and yield sub-optimal solutions.

**Results:** We present a new Integer Linear Programming (ILP) solution for maximum likelihood reconstruction of ancestral PPI networks using the DMC model. We prove the correctness of our solution that is designed to find the optimal solution. It can also use efficient heuristics from general-purpose ILP solvers to obtain multiple optimal and near-optimal solutions that may be useful in many applications. Experiments on synthetic data show that our ILP obtains solutions with higher likelihood than those from previous methods, and is robust to noise and model mismatch. We evaluate our algorithm on two real PPI networks, with proteins from the families of bZIP transcription factors and the Commander complex. On both the networks, solutions from our ILP have higher likelihood and are in better agreement with independent biological evidence from other studies.

**Availability:** A Python implementation is available at https://bitbucket.org/cdal/.

## 1 Introduction

An organism’s genotype and phenotype is mediated by complex biological interactions. Snapshots of such interactions are graphically captured by networks and spatio-temporal analysis of biological networks has led to deep functional and evolutionary understanding of molecular and cellular processes (Yamada and Bork, 2009). Knowledge of the evolution of networks such as Protein-Protein Interactions (PPI), metabolic and gene regulatory networks has been effectively used in the study of: molecular mechanisms in yeast (Wagner, 2001), cell signaling and adhesion genes (Nichols *et al.*, 2006), modularity in metabolic networks of bacterial species (Kreimer *et al.*, 2008), and of protein complexes (Pereira-Leal *et al.*, 2006), functional modules from conserved ancestral protein-protein interactions (Dutkowski and Tiuryn, 2007), evolutionary trends of biosynthetic capacity loss in parasites (Borenstein and Feldman, 2009), regulatory network inference (Zhang and Moret, 2010) and essential and disease-related genes in humans (Vidal *et al.*, 2011).

Generative models, called network growth models, that describe the evolution of networks have been used to explain properties of networks in other domains, such as the Preferential Attachment Model (Barabási and Albert, 1999) (for the World Wide Web) and the Forest Fire Model (Leskovec *et al.*, 2005) (for social networks). These models encode assumptions of evolutionary processes in terms of graph operations. The key evolutionary process characterizing biological networks is duplication and divergence (Wagner, 2001). Thus each evolutionary step is modeled by duplication of a network node (including its incident edges) and deletion of some of the incident edges. Such models have been elucidated and validated in several biological studies (Chung *et al.*, 2003; Vázquez *et al.*, 2003). In this work we use the Duplication-Mutation with Complementarity (DMC) model, that has been found to fit PPI networks better than other commonly used network growth models (Middendorf *et al.*, 2005; Navlakha and Kingsford, 2011).

Similar to reconstruction algorithms to infer evolutionary history of sequences, we can use a network growth model to obtain principled model-based reconstruction of ancestral networks. Assuming such a generative model, ancestral reconstruction seeks to find the most likely sequence of networks that yields the extant network. This entails inferring the order in which nodes duplicate and edges are lost at each step during evolution. Several algorithms have been designed for ancestral network reconstruction. An algorithm for maximum likelihood ancestral reconstruction based on the DMC model, called ReverseDMC, was developed by Navlakha and Kingsford (2011). ReverseDMC greedily (by maximizing the likelihood of that single step) chooses an *anchor* node that is duplicated, at each step of evolution.

ReverseDMC uses only extant network topology to infer ancestral networks. Variants that can use additional biological information of the extant proteins, when available, for ancestral reconstruction have also been proposed. Such additional information include protein duplication history (Li *et al.*, 2013; Jasra *et al.*, 2015) and evolutionary periods of proteins (Zhang *et al.*, 2017). Other techniques for ancestral network reconstruction include the use of graphical models (Pinney *et al.*, 2007), and parsimony-based approaches that find one or more ancestral reconstructions with the minimum number of interaction gain/loss events (Patro *et al.*, 2012; Patro and Kingsford, 2013). These methods also use the gene duplication history and extant networks of multiple species during ancestral network reconstruction. Most of these methods, including ReverseDMC, yield only one evolutionary history, which is obtained by optimizing a mathematical criterion (like likelihood). In many applications it is useful to obtain multiple optimal and near-optimal histories to explore their biological relevance, through alternative criteria.

In this paper, we develop an Integer Linear Programming (ILP) solution for maximum likelihood reconstruction of ancestral PPI networks, using only extant network information. We use indicator variables to determine anchor and duplicated nodes at each step of evolution. Conditions imposed by the DMC model are formulated as linear constraints on each consecutive pair of networks during evolution. We prove the correctness of our algorithm that can find the optimal solution, i.e., a solution that maximizes DMC-model based likelihood.

It is not known whether this problem is polynomial-time solvable. However, it appears to be unlikely, since the number of possible histories grows exponentially with each step. The advantage of an ILP framework is that it can leverage accurate and efficient heuristics, which are being steadily improved by the optimization community with readily available implementations in state-of-the-art general-purpose solvers (Gurobi, 2015). These improvements can automatically enhance the solution quality for the ancestral reconstruction problem. Another advantage of using ILP heuristics is that they can find multiple optimal and near-optimal solutions during their search of the solution space. Thus, they yield multiple reconstructions that can be examined for their biological relevance.

In experiments with synthetic datasets, our ILP solution obtains reconstructions with higher likelihood than those from ReverseDMC, which also shows that the greedy heuristic for this problem is not optimal. Simulation studies also show that our reconstruction algorithm is robust to noise in the extant network and mismatch in input model parameters. We evaluate our algorithm on two real biological networks that contain protein-protein interactions from the families of bZIP transcription factors and the Commander complex. Our ILP obtains solutions with higher likelihood on both these networks. We also examine the biological relevance of the results by comparing the inferred node arrival times as well as the chosen duplicated nodes at each evolutionary step, in reconstructions from ReverseDMC and ILP. On both the networks, solutions from our ILP are in better agreement with independent biological evidence from ortholog information and sequence similarity.

## 2 Problem Statement

Given a network *G*_*t*_ at time *t*, and a model of evolution 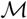 that specifies a series of operations that generates *G*_*g*+1_ from *G*_*g*_, we want to find the most probable sequence of networks *G*_*S*_ = *G*_1_, *G*_2_,…, *G*_*t*−1_:

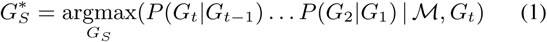

We now describe the model that we use and how likelihood is computed for the model, as given in Navlakha and Kingsford (2011).

The duplication-mutation with complementarity (DMC) model assumes the *G*_2_ to be a simple, connected two–node graph, has two parameters *q*_con_ and *q*_mod_, and network evolution, from any network *G*_*g*_ to *G*_*g*+1_, proceeds as follows (see fig. 1):

**Fig. 1.**
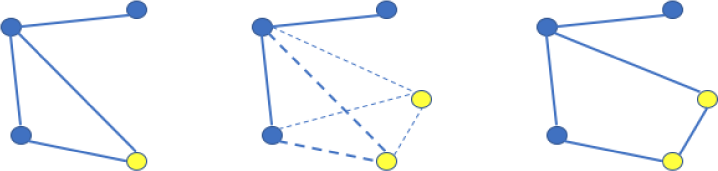
DMC Model. Left: Yellow anchor node selected. Middle: Anchor node is duplicated, with edges to all neighbors. Right: Some edges to neighbors are deleted (with probability *q*_mod_/2), edge between the duplicated nodes retained with probability *q*_con_.

1. An anchor node *u* in *G*_*g*_ is selected at random and duplicated to form node *v*. Initially *v* is connected to all neighbors of *u* and to no other nodes.
2. For each neighbor *x* of *u* (*x* is also a neighbor of *v*), the connecting edge (*u, x*) or (*v, x*) is modified with probability *q*_mod_; if the edge is to be modified, then with equal probability, either edge (*u, x*) or (*v, x*) is deleted.
3. Edge (*u, v*) is added with probability *q*_con_.

Since each time-step adds a node we denote each network by the number of nodes contained in it: *G*_*g*_ is a network with *g* nodes.

Let *e*_*uv*_ denote the edge between the anchor (*u*) and duplicated node (*v*), that is set to 1 if the edge exists and is 0 otherwise. From step 2 of the DMC model, the probability that *u* and *v* share a particular neighbor is (1 − *q*_mod_) and the probability that a node *x* is a neighbor of *u* and not of *v* or a neighbor of *v* and not of *u* is *q*_mod_/2. Let *N*(*u*) denote the neighbors of *u*, the intersection *N* (*u*) *∩ N* (*v*) is the set of common neighbors of *u* and *v* and the symmetric difference *N*(*u*)∆*N*(*v*) is the set of nodes that are neighbors of either *u* or *v* but not both. Then, given *u* and *v* are the anchor and duplicated nodes respectively in *G*_*g*_, we have, ignoring constant terms:

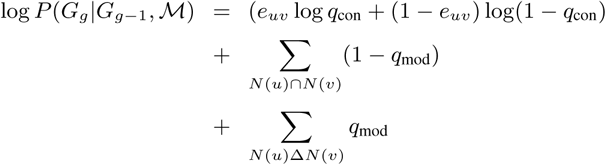

Once *u* and *v* are identified, *G*_*g*−1_ can be reconstructed by removing node *v* and adding edges between *u* and each node in *N*(*u*) − (*N*(*u*) ∩ *N*(*v*)), since these edges were present before step 2 of the DMC model. Note that *u* and *v* are indistinguishable in *G*_*g*_: either one of them may be deleted to form *G*_*g*−1_ and the addition of edges follows *mutatis mutandis*. In the following we will refer to the pair of nodes *u, v* in *G*_*g*_ as *duplicated nodes* and *u* in *G*_*g*−1_ as the *anchor node*.

## 3 ILP-based Solution

### 3.1 Characterizing a Reconstructed Sequence

We first state a theorem that characterizes a sequence of networks that evolves following the DMC model. This forms the basis of the ILP developed in the following section and is also used in proving the correctness of the ILP solution.

Let *G*_*g*_ = {*N*_*g*_, *E*_*g*_} and *G*_*h*_ = {*N*_*h*_, *E*_*h*_} be two networks, with *N, E* representing their sets of nodes and edges, respectively. Nodes and edges in {*N*_*g*_, *E*_*g*_ are identified with the superscript *g* (e.g., node *u*_*g*_ ∈ *N*_*g*_, edge *e*^*h*^ ∈ *E*_*h*_). We use *e*_*φ*_ as an indicator for a ‘dummy edge’ that does not exist in a network. An edge is represented by the pair of nodes it is incident on. For sets *A, B, A/B* denotes the set *A − B*.

#### Definition 3.1.

A pair of networks (*G*_*g*_, *G*_*h*_) is *DMC-evolvable* if *G*_*h*_ can be obtained from *G*_*g*_ through the DMC model of evolution.

#### Theorem 3.1.

A network *G*_*g*_ is *DMC-evolvable* into network *G*_*h*_ iff:

1. |*N*_*h*_| = |*N*_*h*_| + 1
2. ∃ a function *f*_*N*_: *N*_*h*_ → *N*_*g*_ and nodes *a*^*g*^ ∈ *N*_*g*_ and *u*^*h*^, *v*^*h*^ ∈ *N*_*h*_ such that
  a. *f*_*N*_(*u*^*h*^) = *f*_*N*_ (*v*_*h*_) = *a*^*g*^
  b. The restriction of *f*_*N*_: *N*_*h*_/{*u*^*h*^, *v*^*h*^} → *N*_*g*_/{*a*^*g*^} is bijective.
3. The function *f*_*E*_: *E*_*h*_ → *E*_*g*_ ∪ {*e*_*φ*_} given by (i) *f*_*E*_ (*x*^*h*^, *y*^*h*^) = (*f*_*N*_ (*x*^*h*^), *f*_*N*_ (*y*^*h*^)) and (ii) *f*_*E*_ (*u*^*h*^, *v*^*h*^) = *e*_*φ*_, if (*u*^*h*^), *v*^*h*^) ∈ *E*^*h*^ is well-defined and such that
  a. ∀ node *b*^*h*^ that is a neighbor of either *u*^*h*^ or *v*^*h*^ (or both), and *b*_*h*_ ≠ *u*^*h*^, *b*^*h*^ ≠ *v*^*h*^, *f*_*E*_ (*u*^*h*^, *b*^*h*^) = (*a*^*g*^, *f*_*N*_ (*b*^*h*^)) and *f*_*E*_ (*v*^*h*^, *b*^*h*^) = (*a*^*g*^, *f*_*N*_(*b*^*h*^)).
  b. The restriction of 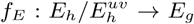 is bijective, where 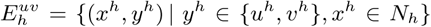, the set of edges incident on *u*^*h*^ and *v*^*h*^.

Proof. (1) follows from the definition of DMC since exactly one node is duplicated at each step. Function *f*_*N*_, *f*_*E*_ define mappings of the nodes and edges respectively, where *a*^*g*^ plays the role of an anchor node in *G*_*g*_ and *u*^*h*^, *v*^*h*^ are the duplicated nodes in *G*_*h*_. Conditions 2(b) and 3(b) ensure that the subgraph in *G*_*g*_ formed by excluding the anchor nodes and its incident edges is isomorphic to the subgraph in *G*_*h*_ formed by excluding the duplicated nodes and their incident edges (see fig. 2). This is true because the DMC model only affects the chosen anchor node and its incident edges, and does not alter any edges or nodes in these subgraphs. The function *f*_*E*_ allows the edge between the duplicated nodes in *G*_*h*_ to remain unmapped to any edge in *G*_*g*_, thereby allowing both cases possible in DMC – there may or may not exist an edge between duplicated nodes. By the DMC model, an incident edge to an anchor node is duplicated and then at most one of the two edges is deleted. This is equivalent to condition 3(a) which ensures that an edge between a duplicated node and a neighbor (that is not the other duplicated node) in *G*_*h*_ is mapped to the anchor node and the mapped neighbor in *G*_*g*_.

Note that the theorem above characterizes DMC-evolvable pairs of networks solely from a combinatorial perspective without taking into account model parameters *q*_*con*_, *q*_*mod*_. The model parameters determine the likelihood of the evolution.

**Fig. 2.**
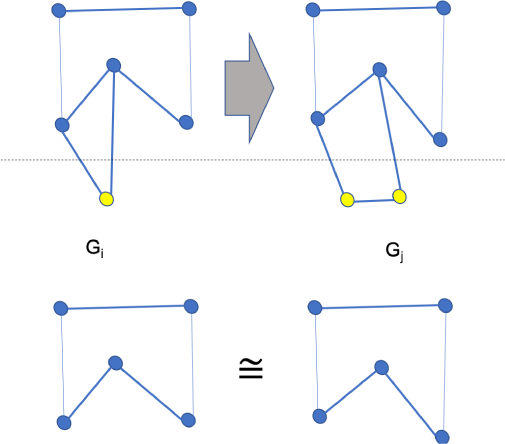
Above: a network evolves following the DMC model. Below: If the anchor node, duplicated nodes and their incident edges are removed, the remaining subgraphs are isomorphic.

#### Definition 3.2.

A sequence of networks *G*_*S*_ = *G*_2_,…, *G*_*t*_ is *DMC-evolvable* if for every pair of consecutive networks (*G*_*i*_, *G*_*i*+1_), *i* = {2, 3,…, *t* − 1}, *G*_*i*_ is *DMC-evolvable* into *G*_*i*+1_.

### 3.2 Our ILP

To recover the entire sequence *G*_*S*_, given the extant network *G*_*t*_, we have to identify the following:

- Anchor nodes in each of the networks *G*_2_,…, *G*_*t*−1_,
- Duplicated nodes in each of the networks *G*_3_,…, *G*_*t*_,
- Edges in each of the networks *G*_3_,…, *G*_*t*−1_.

We will construct an Integer Linear Program (ILP) to obtain the solution. For each graph, *G*_2_,…, *G*_*t*_, we will use binary edge indicators *e*_*ijg*_ that denote presence or absence of an edge and binary node indicators *x*_*ig*_, *y*_*ig*_, *z*_*ig*_, *a*_*ig*_. Subscripts *i, j* refer to nodes and *g* refers to network *G*_*g*_ that has nodes 1,…, *g*. We will set *x*_*ig*_ to 1 if the *i*^*th*^ node in *G*_*g*_ is a duplicated node and *a*_*ig*_ to 1 if the *i*^*th*^ node in *G*_*g*_ is an anchor node. To identify a common neighbor of the duplicated nodes, we will use the indicator *y*_*ig*_ and to identify a neighbor of either one of the duplicated nodes (but not both), we will use the indicator *z*_*ig*_. Note that *e*_*ijg*_, ∀_*i, j*_ are known in networks *G*_2_ and *G*_*t*_ and unknown in all the other networks. All the binary node indicators are unknown in all the networks.

The log of the probability in equation 1 can now be expressed as:

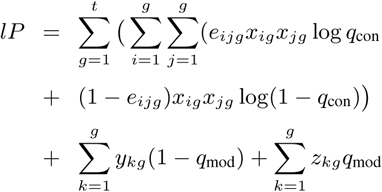

Thus we want to maximize *lP* subject to all the constraints (2 to 23 below) posed by the extant graph and the model, which we shall now describe.

### 3.3 Anchors, Duplicated Nodes and Neighbors

Each network, except *G*_2_, has exactly 2 duplicated nodes:

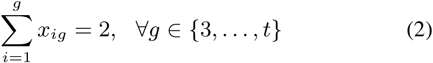

Each network, except *G*_*t*_, has exactly 1 anchor node:

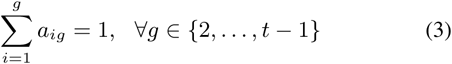

The product *e*_*ijg*_*x*_*ig*_ is 1 if and only if the *i*^*th*^ node is a duplicated node and there is an edge from the *j*^*th*^ node to the *i*^*th*^ node. If the *k*^*th*^ node is a common neighbor there should be exactly 2 edges to the duplicated nodes in the network. Since there are only 2 duplicated nodes per network, for the *k*^*th*^ node, the sum 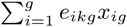 can take only three values: 0,1 or 2. For values 0 and 1, constraint 4 sets *y*_*kg*_ = 0 and for value 2, constraints 4 and 5 set *y*_*kg*_ = 1.

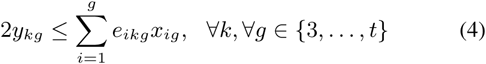

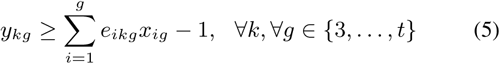

To identify a neighbor of one of the duplicated nodes, but not both, i.e. to set *z*_*kg*_, there should be exactly 1 edge to the duplicated nodes in the network. We also have to ensure that one of the duplicated nodes, which may also satisfy this criterion if the duplicated nodes have an edge between them, is not selected. We can pose these constraints using an auxiliary binary node variable *w*_*kg*_:

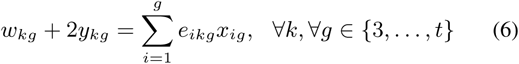

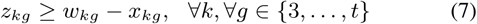

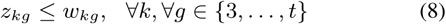

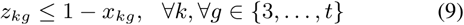

Since there are only 2 duplicated nodes per network, for the *k*^*th*^ node, the sum 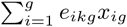 can take only three values: 0,1 or 2.

- If the value is 2, then constraints 4 and 5 ensure that *y*_*kg*_ = 1 and constraint 6 sets *w*_*kg*_ = 0 yielding *z*_*kg*_ = 0 through constraint 8.
- If the value is 1, then *w*_*kg*_ = 1 since constraints 4 and 5 ensure that *y*_*kg*_ = 0. In this case if *x*_*kg*_ = 1 then constraint 9 ensures that *z*_*kg*_ = 0 and if *x*_*kg*_ = 0 then constraint 7 ensures that *z*_*kg*_ = 1.
- Finally, if the value is 0, then *w*_*kg*_ = 0 (constraints 4, 5, 6) and *z*_*kg*_ = 0 through constraint 8.

We use another binary node variable *n*_*kg*_ to indicate a neighbor of a duplicated node, which may be a common neighbor or neighbor of either of the duplicated nodes:

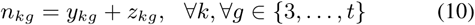

### 3.4 Phantom Edges

During reconstruction, we have to learn the correspondence between nodes in *G*_*g*_ and nodes in the previous network *G*_*g*−1_ to set the values of the unknown edges. In particular, we want to associate the duplicated nodes in network *G*_*g*_ with the anchor node in *G*_*g*−1_. To learn this association, we use indicator variables 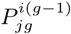 for pairs of nodes (*i*_*g*−1_, *j*_*g*_) where the subscript indicates the network to which the node belongs. Since these are edges that do not exist in the network, but are artificial constructions for our inference, we call them *phantom edges*. We can view them as directed edges to a network from the previous network. See fig. 3 for an illustration.

**Fig. 3.**
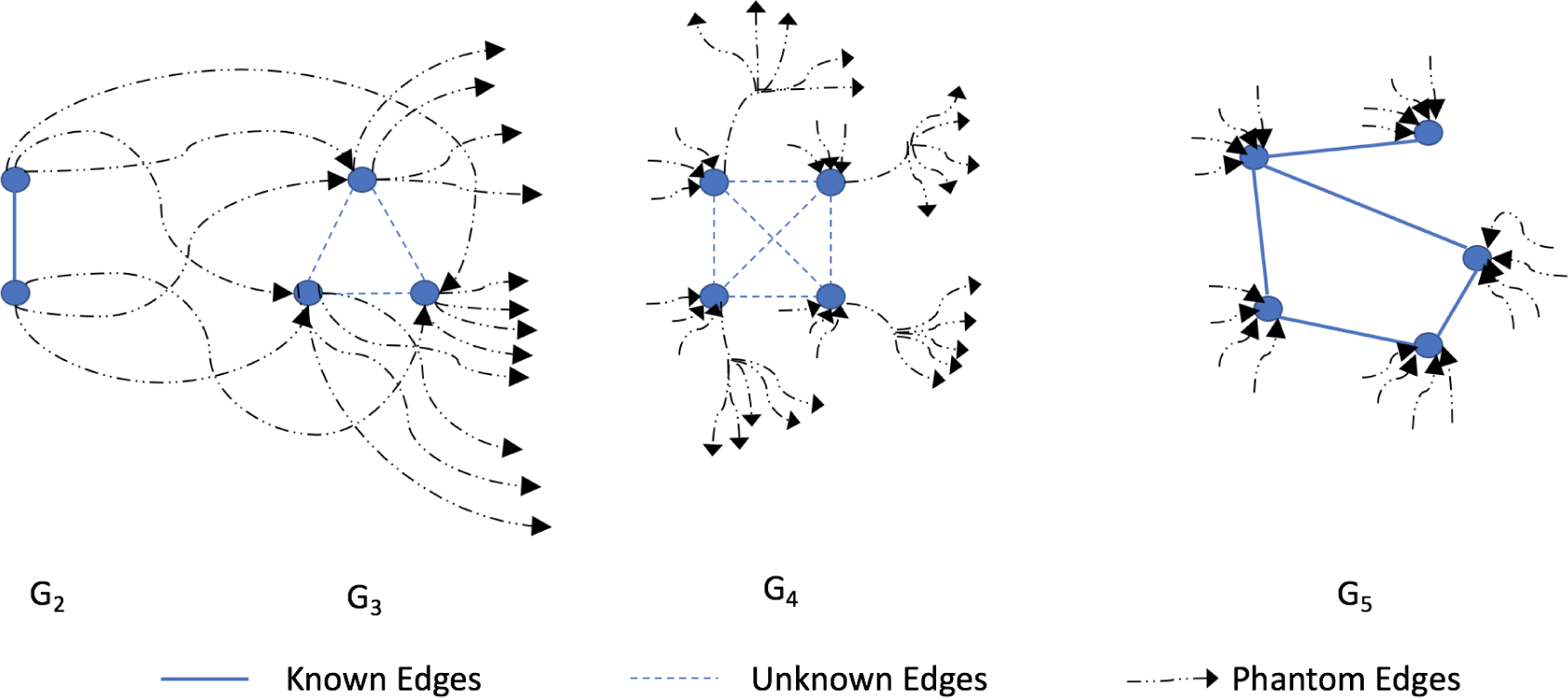
Phantom edges between networks. Each node *i*_*g*_ of a network *G*_*g*_ is connected to all the nodes *j*_*g*_ in *G*_*g*_ through phantom edges 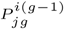. Not all phantom edges shown.

On each node *j*_*g*_ in a network, except in *G*_2_, there must be exactly one incoming phantom edge from any of the nodes (*i*_*g*−1_) in the previous network:

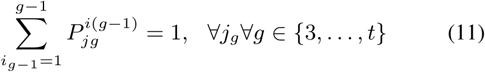

From each node (*i*_*g*−1_) in the (previous) network, except from *G*_*t*_, there must be at least 1 and at most 2 outgoing phantom edges. Anchor nodes will have 2 phantom edges and all other nodes will have only 1:

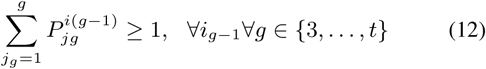

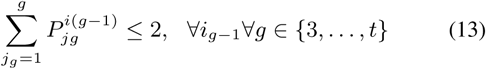

### 3.5 Edge Reconstruction

We now add the final set of constraints for edges in all the ancestral networks that are determined by the model and edges in the extant network. This is done by *mapping* edges from *G*_*g*_ to *G*_*g*−1_ for which we will use the phantom edges. The known edges in the extant network shall be mapped backwards up to the first graph *G*_2_. We have to ensure the following three conditions:

1. An edge between duplicated nodes should not be mapped to any edge in the previous network since the duplicated nodes are from a single anchor node.
2. An edge (*x*_*g*_, *n*_*g*_) between a duplicated node *x*_*g*_ and its neighbor *n*_*g*_ in network *G*_*g*_ should be mapped to an edge (*a*_*g*−1_, *n*_*g*−1_) between the anchor *a*_*g*−1_ and its neighbor *n*_*g*−1_ in network *G*_*g*−1_.
3. Any other edge should be mapped back to a unique edge in the previous network and there should be no other unmapped edge in the previous network.

To set these constraints, we will use three variables defined as follows. A binary indicator variable, for two nodes *i*_*g*_ and *j*_*g*_ in *G*_*g*_, is defined as

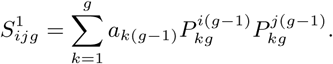

It is non-zero if and only if there are two phantom edges from an anchor node *k*_*g*−1_ in *G*_*g*−1_ to *i*_*g*_ and *j*_*g*_ in *G*_*g*_. For each edge (*i, j*), each term in 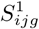 is the product of 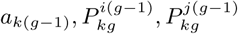. This term has value 1 iff 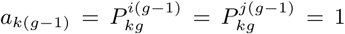 which creates a mapping from nodes *i, j* to the anchor node in the previous network. See fig. 4.

**Fig. 4.**
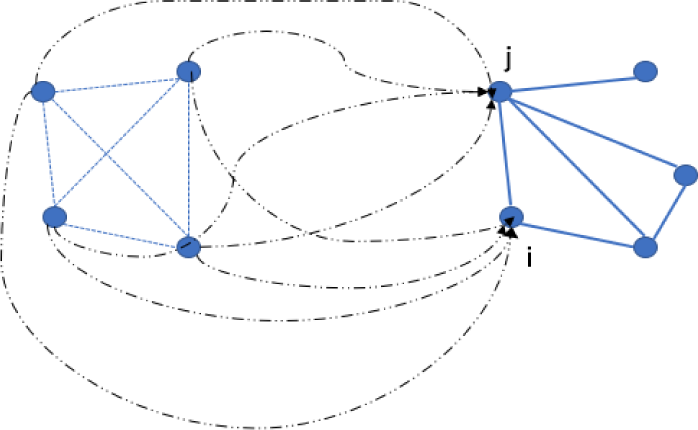
For each pair of nodes (*i, j*), we use phantom edges to find the appropriate mapping. Variables 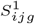, 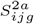, 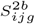, *T*_*ijg*_ encode different possible conditions all of which are not true at the same time. 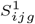 is used to map *i, j* to a single anchor node 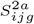, 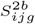 are used to map *i, j* to an anchor node and its neighbor and *T*_*ijg*_ is used for all other cases.

Another binary indicator variable, for two nodes *i*_*g*_ and *j*_*g*_ in *G*_*g*_, is defined as

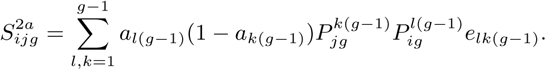

It is non-zero if an only if there are two phantom edges from an anchor node 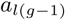 and its neighbor 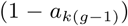 connecting them respectively to *i*_*g*_ and *j*_*g*_ in *G*_*g*_ and there is an edge 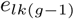 in *G*_*g*−1_. For a symmetric condition, for phantom edges from an anchor node 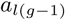 and its neighbor 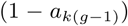 connecting them respectively to *j*_*g*_ and *i*_*g*_ in *G*_*g*_, we define another binary indicator variable, for two nodes *i*_*g*_ and *j*_*g*_ in *G*_*g*_, as

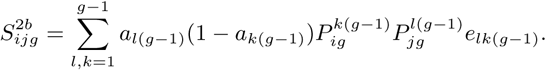

Each term in the sums 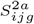 and 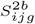 is used to create a mapping from nodes *i, j* to an anchor node and its neighbor in the previous network. See fig. 4.

Finally, another binary indicator variable, for two nodes *i*_*g*_and *j*_*g*_ in *G*_*g*_, is defined as

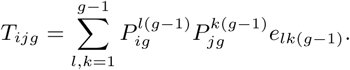

It is non-zero if and only if there are two phantom edges from (any) nodes *k*_*g*−1_ and *l*_*g*−1_ in *G*_*g*−1_ to *i*_*g*_ and *j*_*g*_ in *G*_*g*_ respectively and there is an edge 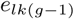. Each term in *T*_*ijg*_ is a product of phantom nodes incoming at *i* and *j* in *G*_*g*_ and the edge *e*_*lk*_(*g*−1) in the previous network *G*_*g*−1_, which when set to 1 creates a mapping from edge (*i, j*) ∈ *G*_*g*_ to edge (*l, k*) ∈ *G*_*g*−1_. See fig. 4.

We set the constraints for each pair of nodes (*i*_*g*_, *j*_*g*_) in graph *G*_*g*_ based on node indicators for duplicated nodes (*x*_*ig*_) and neighbor nodes (*n*_*ig*_):

- If both (*i*_*g*_, *j*_*g*_) are duplicated nodes, i.e. *x*_*ig*_*x*_*jg*_ = 1, then we have to set 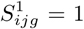 to ensure that duplicated nodes connect to an anchor node in the previous network. Other indicators, 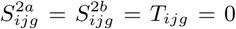 to ensure that no edge in *G*_*g*−1_ is mapped to an edge, if any, between *i*_*g*_ and *j*_*g*_. See fig. 5.
- If the nodes (*i*_*g*_, *j*_*g*_) are such that one of them is a duplicated node and the other a neighbor, i.e. *x*_*ig*_*n*_*ig*_ = 1 or *x*_*jg*_*n*_*ig*_ = 1, then we set 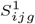 = 0 so the anchor node in the previous network does not connect to this pair through any phantom edges, and we set 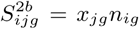, 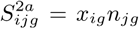 to ensure that phantom edges connect the anchor and its neighbor in the previous graph to nodes (*i*_*g*_, *j*_*g*_). Note that there may not be an edge between (*i*_*g*_, *j*_*g*_), if *j*_*g*_ is a neighbor to the other duplicated node and not *i*_*g*_ as shown in fig. 6. Since both the duplicated nodes map to the anchor, this constraint is set as required. We set *T*_*ijg*_ = 1 to ensure that there is exactly one edge between (*l*_*g*−1_, *k*_*g*−1_) and *T*_*ijg*_ ≥ *e*_*ijg*_ since there may or may not be an edge between (*i*_*g*_, *j*_*g*_). Note that this and the previous cases are mutually exclusive since *n*_*ig*_ and *x*_*ig*_ are never both set to 1 for the same node.
- If both the above cases are not true, i.e. *x*_*ig*_*x*_*jg*_ = *x*_*ig*_*n*_*ig*_ = *x*_*jg*_*n*_*ig*_ = 0, then we set 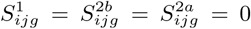 since we do not want an edge between (*i*_*g*_, *j*_*g*_) to map to any edge connecting to an anchor in the previous network and we set *T*_*ig*_ = *e*_*ijg*_ to ensure that there is a single edge (*l*_*g* − 1_, *k*_*g* − 1_) if *e*_*ijg*_ = 1. If *e*_*ijg*_ = 1 then this ensures there is no edge in the previous network mapped to (*i*_*g*_, *j*_*g*_). See fig. 7. The above three sets of conditions are incorporated in the following constraints:

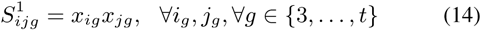

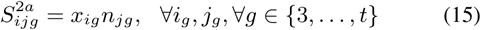

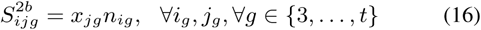 We define an auxiliary binary variable *P*_*ijg*_ that is set to 0 if 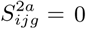 and 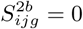 and 1 otherwise (i.e. the logical OR); also, we set *Q*_*ijg*_ = *x*_*ig*_*x*_*jg*_:

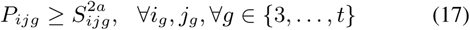

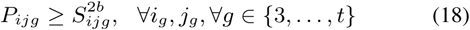

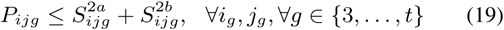 Variables *T*_*ijg*_ and *e*_*ijg*_ are set using *P*_*ijg*_ and *Q*_*ijg*_:

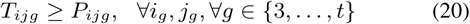

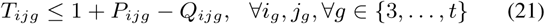

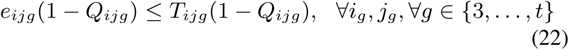

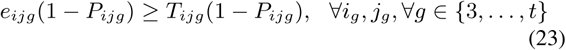
- If *Q*_*ijg*_ = *x*_*ig*_*x*_*jg*_ = 1, then constraints 21 and 22 ensure that *T*_*ijg*_ = *P*_*ijg*_ = 0 since both 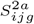 and 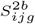 are 0. If *P*_*ijg*_ = 0, *Q*_*ijg*_ = 1, then constraint 22 is void and constraint 23 ensures that *e*_*ijg*_ ≥ *T*_*ijg*_.
- If *Q*_*ijg*_ = *x*_*ig*_*x*_*jg*_= 0 and *P*_*ijg*_ = 1 (i.e. either 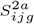 or 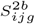 is 1 which is only possible if *x*_*ig*_*n*_*ig*_ = 1 or *x*_*ig*_*n*_*ig*_ = 1) then constraint 20 ensures that *T*_*ijg*_ = 1. If *P*_*ijg*_ = 1, *Q*_*ijg*_ = 0, then constraint 23 is void and constraint 22 ensures that *e*_*ijg*_ ≤ *T*_*ijg*_.
- If *Q*_*ijg*_ = *x*_*ig*_*x*_*jg*_ = 0, and *P*_*ijg*_ = 0, then constraints 20 and 21 do not impose any value on *T*_*ijg*_. If *P*_*ijg*_ = 0, *Q*_*ijg*_ = 0, then constraints 22 and 23 ensure that *e*_*ijg*_ = *T*_*ijg*_.

**Fig. 5.**
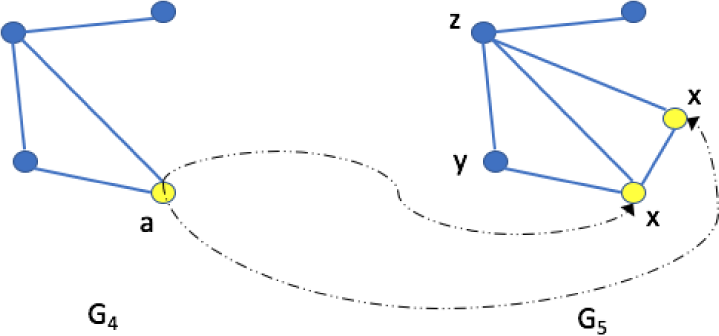
If both (*i*_*g*_, *j*_*g*_) are duplicated nodes (denoted by *x*), then 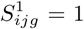 shall connect the duplicated nodes to a single anchor node (denoted by *a*) in the previous network.

**Fig. 6.**
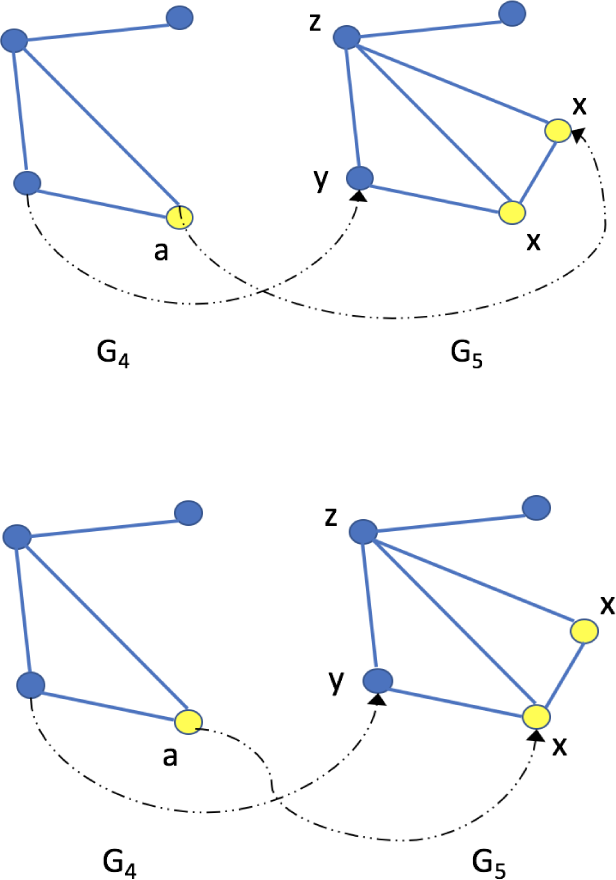
A duplicated node (*x*) and a neighbor (*y* or *z*) must connect to an anchor (*a*) and its neighbor in the previous network. This is done through the variables 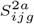, 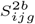. Above: A duplicated node and neighbor of the other duplicated node, Below: A duplicated node and its own neighbor. Both have to be mapped to the same two nodes in the previous network.

**Fig. 7.**
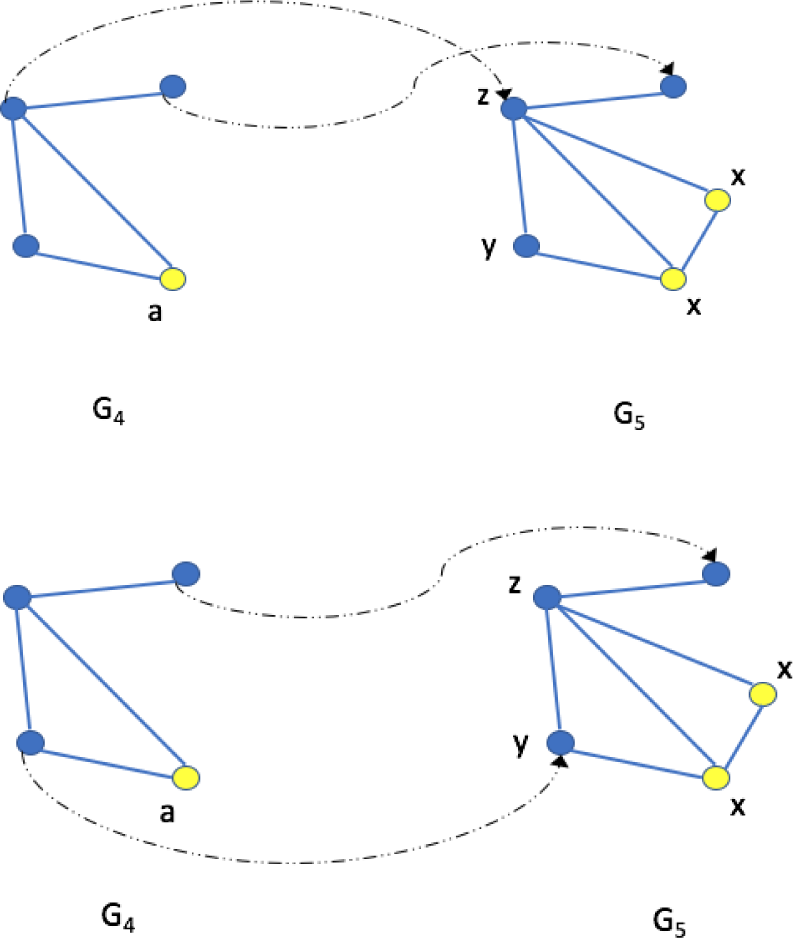
Above: An edge between non-duplicated nodes is mapped back to an edge in the previous network. Below: If there is no edge between a pair of non-duplicated nodes, there should be no edge in the mapped nodes in the previous network.

Finally, we set *e*_*ijg*_ = *e*_*jig*_,*i*_*g*_, *j*_*g*_, ∀_*g*_ ∈ {2,…, *t*} to ensure that the edges are undirected.

### 3.6 Proof of Correctness

#### Theorem 3.2.

A feasible solution to the above ILP yields a DMC-evolvable sequence of networks.

Proof. We use the conditions stated in theorem 3.1. Let *G*_*g*_ = (*N*_*g*_, *E*_*g*_) and *G*_*g*−1_ = (*N*_*g*−1_, *E*_*g*−1_) be any two consecutive networks in the ILP solution. We show that they are DMC-evolvable. Condition (1) is true by construction. The anchor nodes *a*^(*g−*1)^ are identified using constraint 3. Similarly the duplicated nodes *u*^*g*^, *v*^*g*^ are identified using constraint 2.

The phantom edges that are set to 1 explicitly define the function *f*_*N*_: *N*_*g*_ → *N*_*g*−1_ (the direction in fig. 11 is that of the pre-image). Constraint 11 ensures that each element of the domain is mapped. Together, constraints 11, 12 and 13 ensure that there will be exactly as many mappings as the number of nodes in *G*_*g*_. Constraint 3 ensures that *G*_*g*_ has exactly one anchor node and constraint 2 ensures that *G*_*g*−1_ has exactly two duplicated nodes. The duplicated nodes are mapped to the previous network’s anchor node, otherwise constraint 14 is violated. This satisfies condition 2(a). Further this also implies that the restriction of 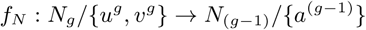 has exactly *|N*_(*g−*1)_*|−*1 mappings. If 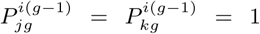 for the same node *i* ∈ *G*_*g* − 1_, then *j* = *k*, otherwise an element in the co-domain is left unmapped which violates constraint 12. Hence, *f*_*N*_ is injective. Constraint 12 ensures that each element of the codomain is mapped and thus *f*_*N*_ is surjective. This proves bijectivity (condition 2(b)).

If there is an edge (*i*^*g*^, *j*^*g*^) between duplicated nodes in *G*_*g*_, i.e., *e*_*ijg*_ = 1 then constraints 21 and 22 ensure that 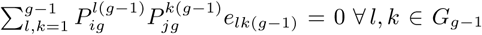. Since *f*_*N*_ is bijective, this is possible only if there is no edge *e*_*lk(g-1)*_ ∈ *G*_*g*-1_ to which this edge is mapped. Thus, the artificial mapping to *e*_*ϕ*_ is enforced for an edge between duplicated nodes, if present. Constraints 15 and 16 directly ensure condition 3(a).

Consider the restriction of 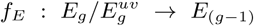. To prove surjectivity, assume the contrary: let an edge in *E*_(*g−1*)_, 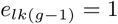 have no preimage, i.e., there is no edge (*i*^*g*^, *j*^*g*^) such that 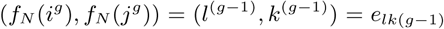. Since *f*_*N*_ is bijective, 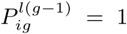 for some *i*^*g*^ and 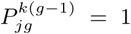 for some *j*^*g*^, and so the product 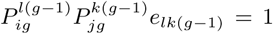 and hence we have *T*_*ig*_ ≥ 1. Note that nodes *i*^*g*^, *j*^*g*^ are not duplicated nodes, since edges incident on them are not in 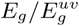, this sets *Q*_*ijg*_ = *P*_*ijg*_ = 0. Thus constraint 23 becomes *e*_*ijg*_ ≥ *T*_*ijg*_. If there is no edge (*i*^*g*^, *j*^*g*^), then *e*_*ijg*_ = 0, and constraint 23 is violated. This shows that *f*_*E*_ is surjective. For an edge 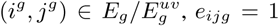, *e*_*ijg*_ = 1 and constraints 22 and 23 imply that 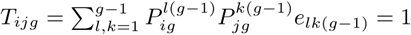 which implies that for exactly one edge 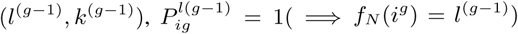 and 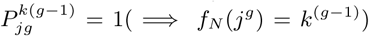 and 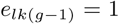 which proves injectivity. Hence *f*_*E*_ is bijective and condition 3(b) is satisfied.

Since any two consecutive networks in the sequence of networks are DMC-evolvable, the reconstructed sequence is DMC-evolvable.

### 3.7 Linearization

The constraints as described above have terms that are products of binary variables and sums of such products. A constraint *y* = *x*_1_*x*_2_,…, *x*_*n*_ where each variable is binary is equivalent to the following *n* + 1 constraints: *y ≥ x*_1_, *y ≥ x*_2_,… *y ≥ x*_*n*_, *y ≤ x*_1_ +*x*_2_ +*…*+*x*_*n*_ (*n* − 1). Sums of products can be decomposed using auxiliary binary variables. For example, *y* = *x*_1_*x*_2_ + *x*_3_*x*_4_ can be expressed as *y* = *z*_1_ + *z*_2_, *z*_1_ = *x*_1_*x*_2_, *z*_2_ = *x*_3_*x*_4_ and further linearized using the previous rule.

### 3.8 Computational Complexity and Multiple Solutions

Since ILP is, in general, NP-hard, optimal solutions for very large networks may not be found in polynomial time. Typically, the worst-case time complexity is exponential in the number of variables. In our formulation, for an extant network of *n* nodes, there are *O*(*n*^2^) node indicator variables, *O*(*n*^3^) edge indicator variables and *O*(*n*^3^) indicator variables for phantom edges.

Many heuristics such as LP-based Branch and Bound and Cutting Plane methods have been developed to find multiple near-optimal solutions (e.g., see (Wolsey, 1998)), with efficient software implementations (Gurobi, 2015). These heuristics are designed to run for a pre-specified period of time, during which they find multiple (near-optimal) solutions. Thus, there is a trade-off between the running time and quality of solutions found.

## 4 Experiments

### 4.1 Simulations

We test the performance of our algorithm and ReverseDMC, the Greedy approach of Navlakha and Kingsford (2011), in simulations where the complete evolutionary history is known. Evolution is simulated following the DMC model starting from an initial network of two connected nodes. As detailed in the following sections, we evaluate three settings:

- Reconstruction from the extant network using the true model parameters *q*_*con*_ and *q*_*mod*_.
- Reconstruction from a noisy extant network using the true model parameters.
- Reconstruction from the extant network when the true model parameters are unknown.

We use three evaluation metrics to assess the reconstructed histories. The likelihood of the entire reconstruction is our first metric. The second metric evaluates the node arrival order during evolution. For the inferred histories this is determined by inverting the list of *removed nodes* from each step of the reconstruction. The correlation between this ranked list and the true arrival order are compared using Kendall’s Tau (Kendall, 1945) (definition given in appendix). Our third metric compares the networks generated at each step of reconstruction with their corresponding true networks. We use graph kernels, which provides a measure of structural similarity between graphs (Vishwanathan *et al.*, 2010). For an extant network *G*_*t*_, given the true sequence of graphs, *G*_*S*_ = *G*_3_,…, *G*_*t*−1_ and a reconstructed history, 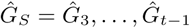, we compute the kernel similarity: 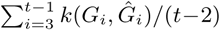, where *k* is a graph kernel. This measures the average similarity of the entire reconstruction. We use the Weisfeiler-Lehman kernel that measures similarity based on isomorphism of subgraphs within the input graphs (Shervashidze *et al.*, 2011). In all three metrics, higher values indicate better performance.

#### Reconstruction with true model parameters

We simulated 1500 extant networks with number of nodes in the extant network varying from 6 to 10. For each simulation, the value of each DMC parameter (*q*_*con*_ and *q*_*mod*_) was randomly chosen from the interval [0.1, 0.9], rounded to one decimal. The same parameters were used during reconstruction, for both ReverseDMC and our ILP.

Figure 8 shows that there were no simulations where solutions from ILP have a lower likelihood than that of ReverseDMC. Since these are relatively small networks, both ReverseDMC and ILP were able to find optimal solutions in 76% of the cases, while in 24% of the cases ILP found solutions with higher likelihood. The fact that ILP could find solutions with higher likelihood shows that ReverseDMC is not guaranteed to find optimal solutions. Histories reconstructed from ILP had better correlation with the true histories, with respect to node arrival order, in 93% of the cases. The kernel similarity values of the reconstructed networks were not lower than those from ReverseDMC in 79% of the cases. Figure 8 also shows the summary statistics of the log-likelihood, Kendall’s Tau and Kernel similarity values. Overall, reconstructed histories from ILP have higher likelihood and obtain node arrival orders and inferred networks that are closer to the true evolutionary history compared to those from ReverseDMC.

**Fig. 8.**
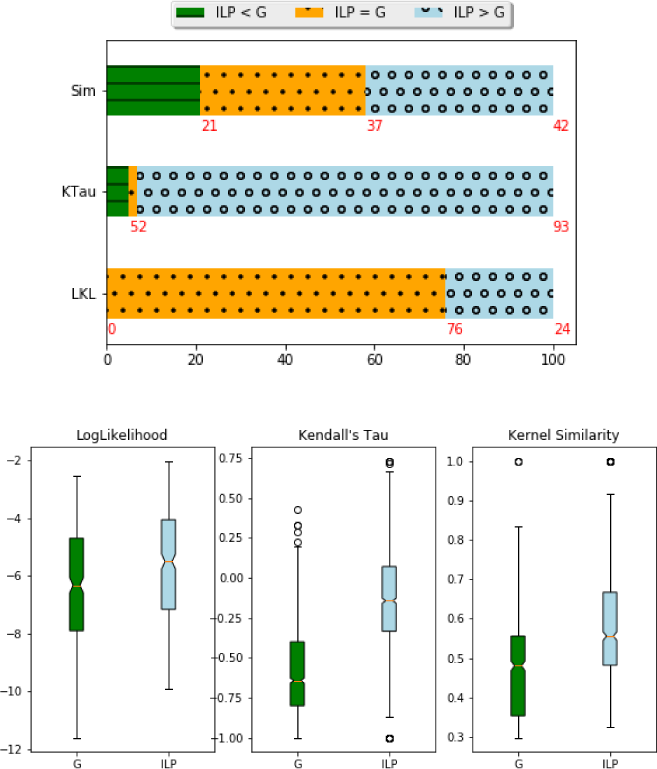
Reconstruction with true model parameters. Above: Proportion of simulations where reconstructed histories from ILP scored higher (ILP > G), equal (ILP = G) and lower (ILP < G) than the reconstructions from ReverseDMC, the Greedy approach of Navlakha and Kingsford (2011), for three metrics - LogLikelihood (LKL), Kendall’s Tau (KTau) and Kernel Similarity (Sim). Numbers below each bar indicate percentages. Below: Summary Statistics of LogLikelihood, Kendall’s Tau and Kernel Similarity values of reconstructions from ILP and ReverseDMC (G).

#### Reconstruction with noisy extant networks

To assess the performance when there are deviations from the growth model, we use extant networks (from our previous experiment, above) and randomly replace *p*% of the edges with new edges. These *noisy* networks are given as input to the reconstruction algorithms. We study the performance of the algorithms at *p* = 10%, 20%.

Figure 9 shows summary statistics of log-likelihood, Kendall’s Tau and Kernel similarity values obtained in our experiments. The likelihood and kernel similarity of the reconstructed solutions from both the algorithms deteriorate with increasing noise in the extant networks. The decrease in the Kendall’s Tau values is lesser, although the number of instances with high values for both algorithms decrease with increasing noise.

**Fig. 9.**
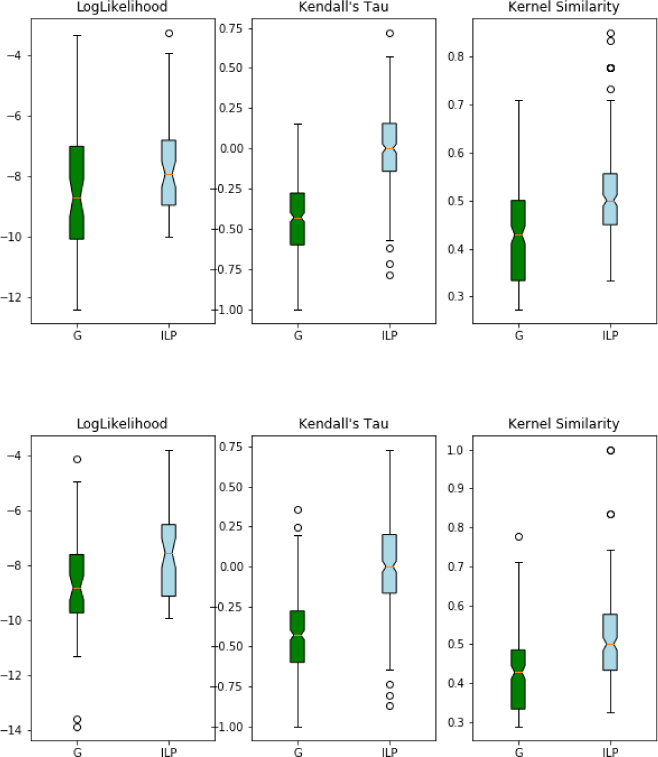
Reconstruction with noisy extant networks. Summary Statistics of LogLikelihood, Kendall’s Tau and Kernel Similarity values of reconstructions from ILP and ReverseDMC at varying noise levels in extant networks (Above: 10%, Below: 20%).

The relative performance trend remains the same as before: reconstructed histories from ILP have higher likelihood and obtain node arrival orders and inferred networks that are closer to the true evolutionary history compared to those from ReverseDMC. This also seen in figure 10 which shows the proportion of solutions where ILP obtains better, equal or worse solutions compared to ReverseDMC with respect to each of the three metrics. There is no significant change in these proportions with increasing noise in the extant networks.

**Fig. 10.**
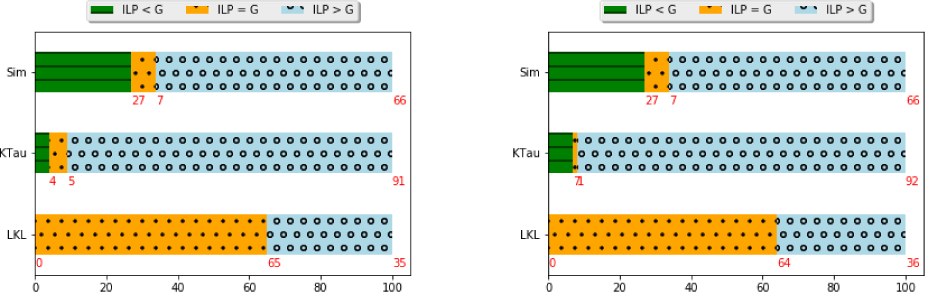
Reconstruction with noisy extant networks. Proportion of simulations where reconstructed histories from ILP scored higher (ILP > G), equal (ILP = G) and lower (ILP < G) than the reconstructions from ReverseDMC, the Greedy approach of Navlakha and Kingsford (2011), for three metrics - LogLikelihood (LKL), Kendall’s Tau (KTau) and Kernel Similarity (Sim). Noise levels: 10% (left), 20% (right).

**Fig. 11.**
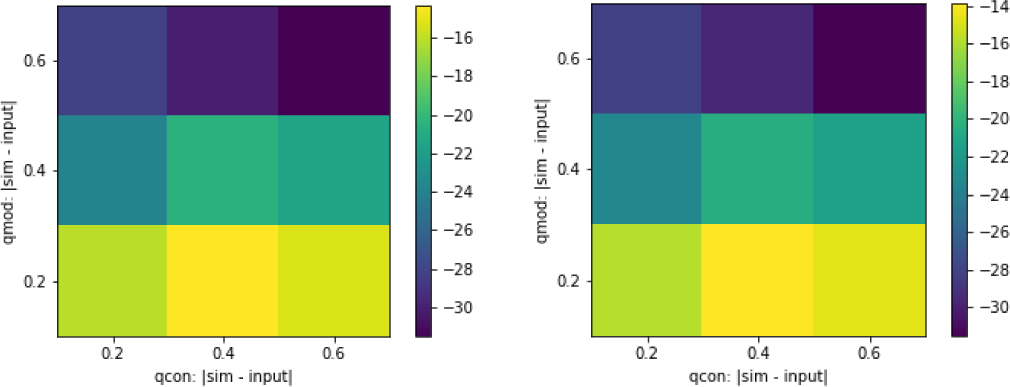
Likelihood of the reconstructed solutions from ReverseDMC (left) and ILP (right). Along the axes: difference between the input parameters to the reconstruction algorithm and the parameter used in simulating the network, for *q*_*con*_ (X-axis) and *q*_*mod*_ (Y-axis).

#### Reconstruction when true model parameters are unknown

We evaluate the performance of the algorithms when the parameters used during reconstruction differ from the model parameters used during simulation. We test this for three sets of simulations where model parameters (*q*_*mod*_, *q*_*con*_) used during evolution were {(0.9,0.1), (0.4,0.4), (0.8,0.2)}. For each case, we provided different valid parameters (*q*_*mod*_ ± ∆, *q*_*con*_ ± ∆) to ReverseDMC and ILP with ∆ = {0.2, 0.4, 0.6}.

Figure 11 shows the likelihood of the reconstructed solutions from ReverseDMC and ILP. In both cases, the performance deteriorates with increasing ∆ and is more affected by mismatch in *q*_*mod*_ values. This is consistent with the findings in Navlakha and Kingsford (2011) for ReverseDMC. Although ILP obtains solutions with higher likelihood (and Kendall’s Tau, Kernel similarity values) overall, the general trend remains the same.

### 4.2 Real Networks

We reconstruct the history of two protein-protein interaction networks using both ReverseDMC and ILP algorithms. Each algorithm is run for a range of values of *q*_*con*_ and *q*_*mod*_, the solution with the best likelihood is chosen for further analysis.

Since the true evolutionary histories are not known, we cannot use the evaluation metrics that we used for our simulation studies. We evaluate the biological relevance of the results in two ways. First, we compare the node arrival times of the reconstructions following the procedure described in Navlakha and Kingsford (2011). The key idea is to estimate the protein arrival time using available ortholog information, with the assumption that proteins that arrive earlier in history have higher number of orthologs. Thus, the list of proteins in the extant network in descending order of number of orthologs is considered to be the ‘true’ node arrival order (*A*_*T*_). We determine the number of orthologs for each protein using OrthoDB (Kriventseva *et al.*, 2018), by counting the number of genes at the highest level at which ortholog information was available for all the proteins in the networks (vertebrata for bZIP and metazoa for Commander). The reconstruction history of both Greedy and ILP identifies the removed node at each step: this provides the reconstructed node arrival order (*A*_*R*_) for each algorithm. *A*_*T*_ and *A*_*R*_ are compared using Kendall’s Tau (Kendall, 1945) that measures correlation between two ranked lists (definition given in appendix). Higher values indicate better correlation.

Our second evaluation is based on the sequence similarities between all the inferred anchors and duplicated nodes. Since at each time step in evolution (by the DMC model) the anchor gene (*a*) duplicates into another gene (*d*), we expect the pairwise similarity between *a* and *d* to be higher than the pairwise similarity between *a* and the remaining genes at that time step. Given the the extant network *G*_*t*_ and its reconstructed evolutionary history: 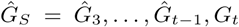, along with chosen anchors and duplicated nodes in each network, we compute a score 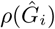 for each network in 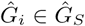, using pairwise sequence similarity (Needleman and Wunsch, 1970) between the chosen anchor node protein and the duplicated node protein. The final score for the reconstruction, that we call Anchor-Duplicate Similarity Score (ADSS), is given by 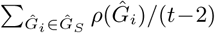, where we normalize by the number of networks in 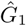. 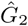 is not considered since in the first evolutionary step (from 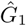 to 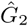 there is only one gene that duplicates and there are no other genes to compare with. Thus given two reconstructions of the same extant network, higher ADSS indicates better choice of anchor and duplicate nodes in the reconstruction.

#### bZIP Transcription Factors

The basic-region leucine zipper (bZIP) transcription factors are a protein family involved in many cellular processes including the regulation of development, metabolism, circadian rhythm, and response to stress and radiation (Amoutzias *et al.*, 2006; Pinney *et al.*, 2007). The interactions between these proteins are strongly mediated by their coiled-coil leucine zipper domains and so, the strength of these interactions can be accurately predicted using just sequence information (Fong *et al.*, 2004). With the method of Fong *et al.* (2004), Pinney *et al.* (2007) constructed extant networks on a set of bZIP proteins for multiple species. We took the *H. sapiens* network and merged subunits for the same protein into one node, to obtain the extant network used in our experiment (fig. 12).

**Fig. 12.**
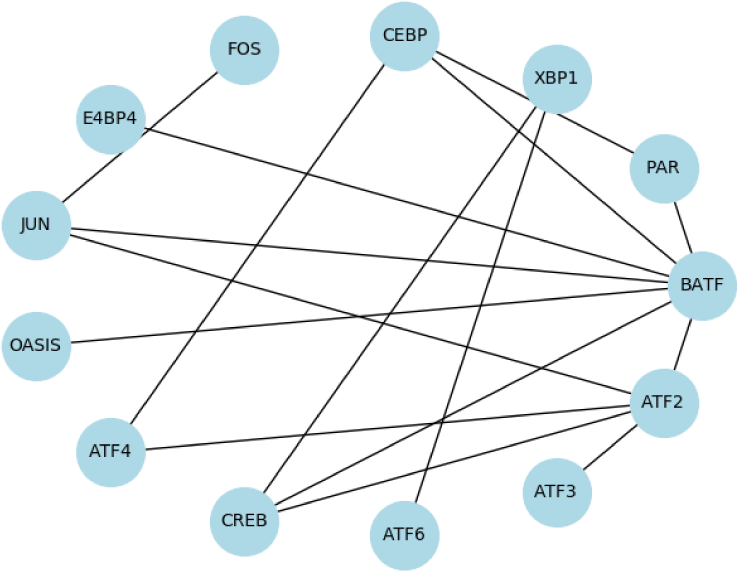
Extant bZIP network used in our experiment.

Table 1 shows the Likelihood, Node Arrival Time Accuracy (measured by Kendall’s Tau) and ADSS for Ancestral Reconstruction of the bZIP Network by both ReverseDMC and ILP. With respect to all three metrics, the solution obtained by ILP is better than that of ReverseDMC. Table 2 shows the order of arrival of proteins inferred by the reconstructions from ReverseDMC and ILP. Sequence-based phylogenetic analysis of bZIP transcription factors by Amoutzias *et al.* (2006) revealed a highly conserved ancient core network containing proteins JUN, FOS and ATF3, that provides additional evidence of the correctness of our reconstruction. In table 2 we observe that these three proteins appear early in the order inferred by ILP (before the seventh step) while JUN and ATF3 arrive after the seventh step in the order inferred by ReverseDMC.

**Table 1.** Likelihood, Node Arrival Time Accuracy and ADSS for Ancestral Reconstruction of the bZIP Network

| Algorithm | Log-Likelihood | Kendall’s Tau | ADSS |
| --- | --- | --- | --- |
| ReverseDMC | -21 | -0.23 | -1901.55 |
| ILP | -19.6 | 0.26 | -1797.11 |

**Table 2.** Arrival order of anchor proteins in the bZIP network, at each step of evolution, based on reconstructions from ReverseDMC and ILP.

| Time step | ReverseDMC | ILP |
| --- | --- | --- |
| 2 | FOS, CREB | ATF2, ATF6 |
| 3 | ATF6 | FOS |
| 4 | BATF | ATF3 |
| 5 | ATF4 | CREB |
| 6 | ATF2 | JUN |
| 7 | E4BP4 | CEBP |
| 8 | JUN | BATF |
| 9 | ATF3 | PAR |
| 10 | CEBP | ATF4 |
| 11 | PAR | E4BP4 |
| 12 | XBP1 | OASIS |
| 13 | OASIS | XBP1 |

#### Commander Network

Commander is a multiprotein complex that is broadly conserved across vertebrates and is involved in several roles including pro-inflammatory signaling and vertebrate embryogenesis (Mallam and Marcotte, 2017). A well characterized sub-complex of Commander, CCC, made of COMMD1, CCDC22, CCDC93 and C16orf62, is known to be involved in endosomal protein trafficking (Bartuzi *et al.*, 2016; Mallam and Marcotte, 2017). Defects in the Commander complex are associated with developmental disorders (Mallam and Marcotte, 2017; Liebeskind *et al.*, 2019). Reconstructing the evolutionary history of interactions in the complex can shed light on the conservation and stability of the proteins and their interactions, which in turn can aid understanding of the sources of dysfunction of the complex. We use the network discussed in Liebeskind *et al.* (2019), shown in fig. 13, as the extant network for ancestral reconstruction.

**Fig. 13.**
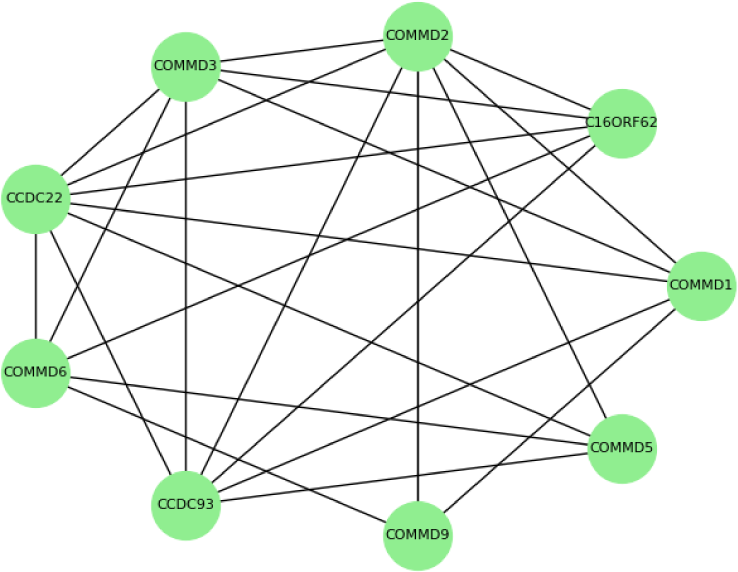
Extant Commander network used in our experiment.

Table 3 shows the Likelihood, Node Arrival Time Accuracy (measured by Kendall’s Tau), and ADSS for ancestral reconstruction by both ReverseDMC and ILP. On this network too, on all three metrics, the solution obtained by ILP is better than that of ReverseDMC. Table 4 shows the order of arrival of proteins inferred by the reconstructions from ReverseDMC and ILP. Among all the commander proteins, COMMD1 is the best studied and is found to be highly conserved with multiple key functions (Riera-Romo, 2018). Indeed, in OrthoDB, COMMD1 has the maximum number of orthologs, among these proteins. In the reconstruction by ILP, COMMD1 is seen to arrive early, at the third step, while in the reconstruction from ReverseDMC it arrives only at the eighth step of evolution.

**Table 3.** Likelihood, Node Arrival Time Accuracy and ADSS for Ancestral Reconstruction of the Commander Network

| Algorithm | Log-Likelihood | Kendall’s Tau | ADSS |
| --- | --- | --- | --- |
| ReverseDMC | -17.49 | -0.27 | -2873.25 |
| ILP | -16.17 | 0.01 | -1649.64 |

**Table 4.** Arrival order of anchor proteins in the commander network, at each step of evolution, based on reconstructions from ReverseDMC and ILP.

| Time step | ReverseDMC | ILP |
| --- | --- | --- |
| 2 | COMMD6, COMMD9 | COMMD3, COMMD9 |
| 3 | C16ORF62 | COMMD1 |
| 4 | COMMD2 | COMMD6 |
| 5 | CCDC93 | COMMD2 |
| 6 | CCDC22 | COMMD5 |
| 7 | COMMD5 | CCDC93 |
| 8 | COMMD1 | CCDC22 |
| 9 | COMMD3 | C16ORF62 |

## 5 Conclusion

We presented an Integer Linear Programming (ILP) based solution for maximum-likelihood reconstruction of the evolution of a PPI network using the Duplication-Mutation with Complementarity (DMC) model. We proved the correctness of the ILP that is designed to find the optimal solution and heuristics from general-purpose ILP solvers can be used to find multiple optimal and near-optimal solutions.

We compared the solutions obtained by our ILP with those from ReverseDMC (Navlakha and Kingsford, 2011), the previous best algorithm for this problem. On simulated data, we found that ILP always obtains solutions that are of equal or higher likelihood than those from ReverseDMC. Further, the ILP solutions are in better agreement with the true evolutionary histories and are robust to noise in extant networks and mismatch in input model parameters. We evaluated both the algorithms on two real PPI networks, containing proteins from the bZIP transcription factors and Commander complex respectively. On both the networks, solutions from our ILP had higher likelihood and were in better agreement with independent biological evidence from ortholog information and sequence similarity.

This is the first ILP solution to a model-based network reconstruction problem and the presented framework may be useful for other network models as well. The ILP framework could be generalized to handle multiple input networks as well as to take into account additional information, such as gene duplication histories. It can also be used as a sub-routine in a larger heuristics. A limitation of our solution is the running time of ILP heuristics – it can take a considerably long time to find good solutions for large networks. However, the ILP framework can yield deeper insights into the structure of the problem and efficient and scalable heuristics could be developed in the future.

## Acknowledgements

Part of the work was performed during XZ’s visit at the Simons Institute for the Theory of Computing at University of California, Berkeley. We thank Rob Patro for sharing the bZIP network data used in their publication (Patro and Kingsford, 2013) which is originally from (Pinney *et al.*, 2007).

## Appendix Kendall’s Tau

We use the version that accounts for ties, given by 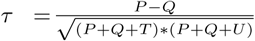, where *P* is the number of concordant pairs, *Q* the number of discordant pairs, *T* the number of ties only in *A*_*T*_, and *U* the number of ties only in *A*_*R*_. If a tie occurs for the same pair in both *A*_*T*_ and *A*_*R*_, it is not added to either *T* or *U*. Here we consider pairs of observatios (*x*_*i*_, *y*_*i*_), (*x*_*j*_, *y*_*j*_) where *x*_*i*_, *x*_*j*_ ∈ *A*_*T*_, *y*_*i*_, *y*_*j*_ ∈ *A*_*R*_ and *i* < *j*. A pair (*x*_*i*_, *y*_*i*_), (*x*_*j*_, *y*_*j*_) is concordant if the ranks of both elements agree, i.e., both *x*_*i*_ < *x*_*j*_ and *y*_*i*_ < *y*_*j*_; or both *x*_*i*_ > *x*_*j*_ and *y*_*i*_ > *y*_*j*_. A pair (*x*_*i*_, *y*_*i*_), (*x*_*j*_, *y*_*j*_) is discordant if *x*_*i*_ > *x*_*j*_ and *y*_*i*_ < *y*_*j*_ or if *x*_*i*_ < *x*_*j*_ and *y*_*i*_ > *y*_*j*_. If *x*_*i*_ = *x*_*j*_ or *y*_*i*_ = *y*_*j*_, it is considered a tie.

